## Supplementary Information for "Conformational Dynamics and Membrane Insertion Mechanism of B4GALNT1 in Ganglioside Synthesis"

**This file contains:**

Supplementary Figures S1-S9

Supplementary Table S1

Supplementary References

**Additional file:**

Supplementary Movie M1

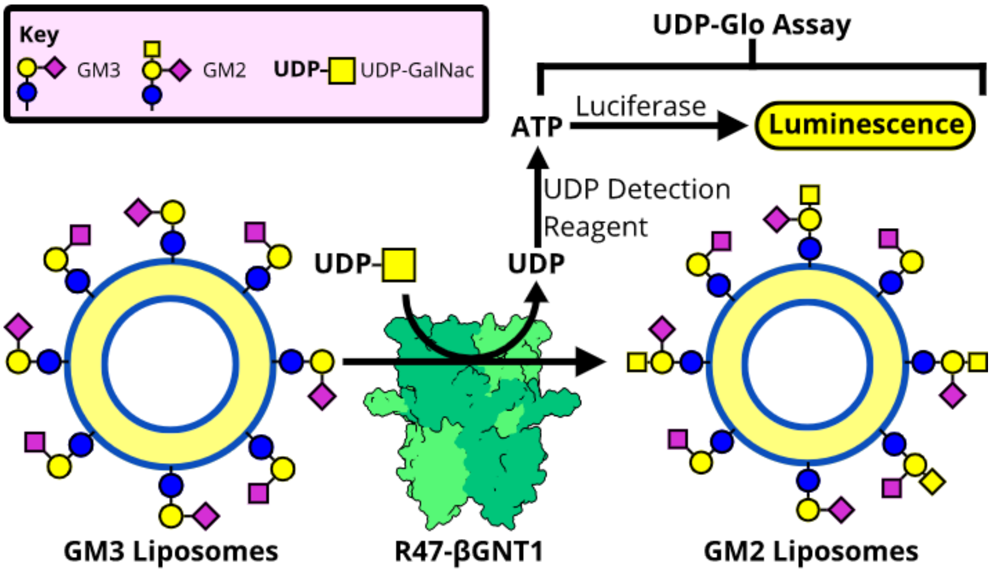


**Supplementary Figure S1. Schematic diagram of the two-step B4GALNT1 activity assay.** The UDP produced by B4GALNT1 following transfer of the GalNac group from UDP-GalNac donor substrate to GM3 lipid substrate is quantified using a UDP-Glo assay. Liberated UDP is converted to ATP and used as a substrate for luciferase producing a product detectable by luminescence.

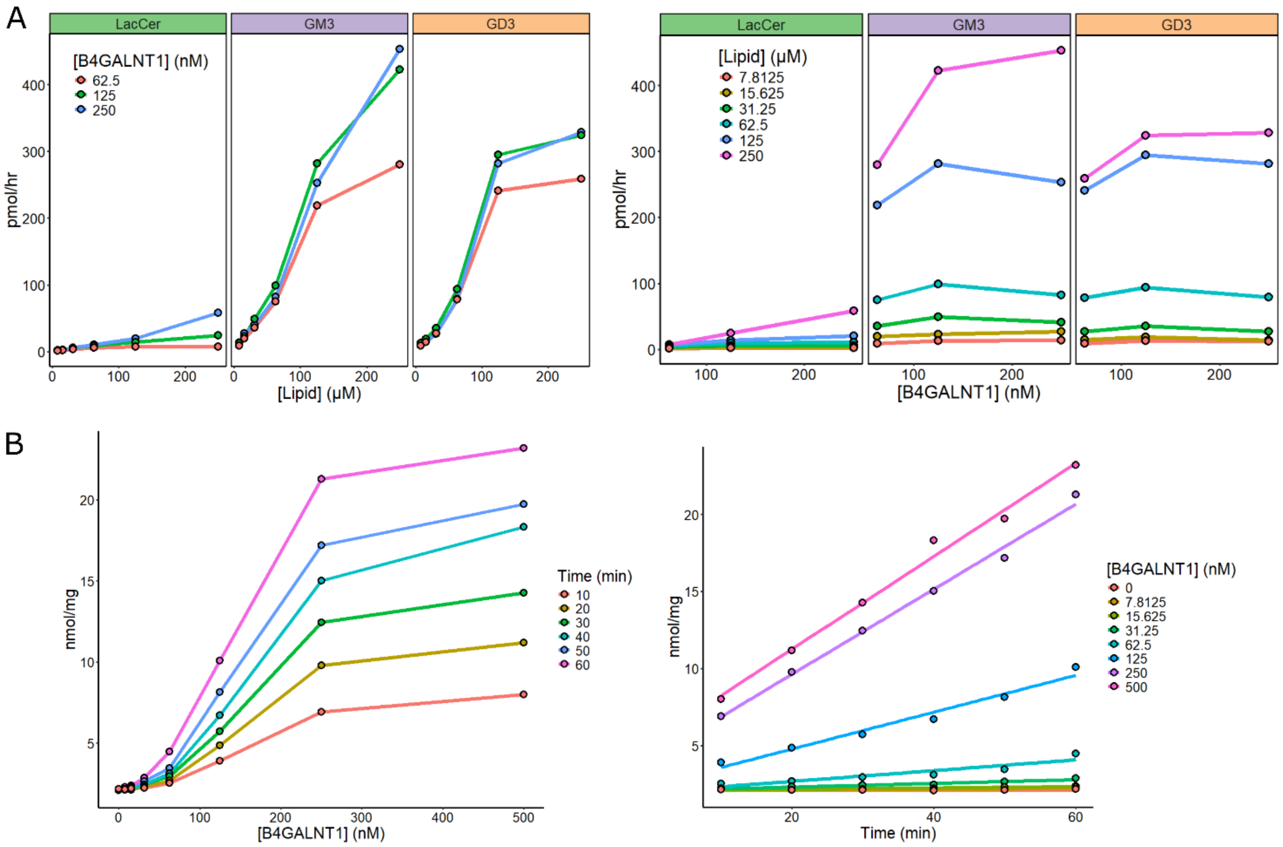


**Supplementary Figure S2. Optimisation of B4GALNT1 activity assay conditions. (A)** B4GALNT1 activity was monitored against the three lipid substrates LacCer, GM3 and GD3 across a range of substrate and enzyme concentrations (n=1). **(B)** Linearity of enzyme activity was tested across a range of enzyme concentrations and over time with ganglioside GM3 (n=1).

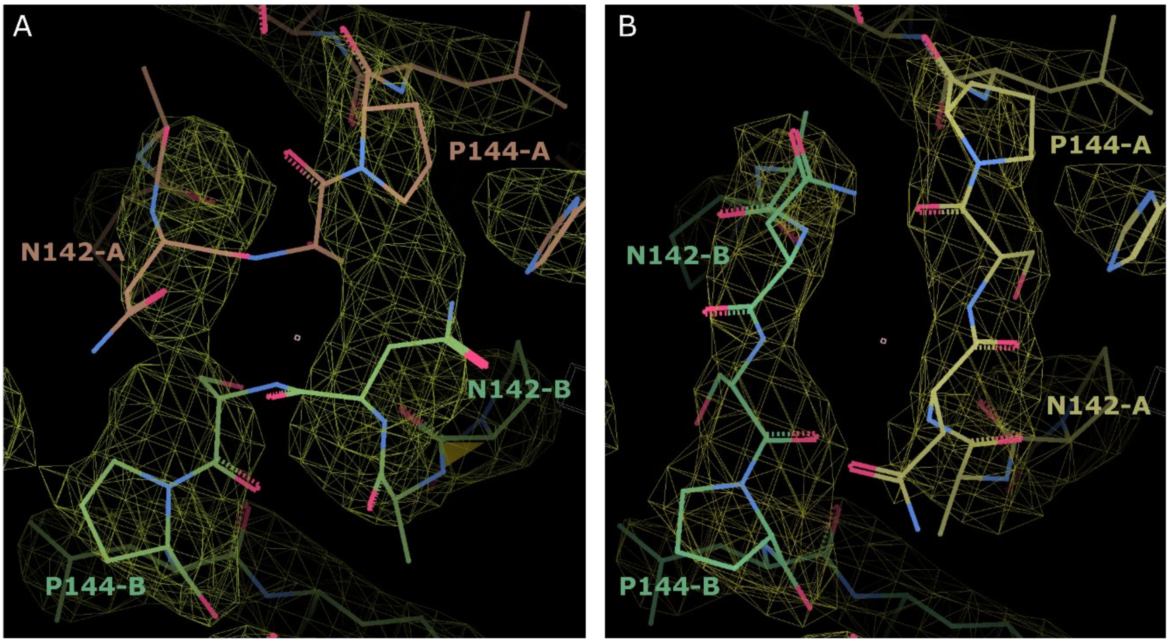


**Supplementary Figure S3. Electron density maps demonstrating rethreading of the B4GALNT1 mainchain in the AF2 model versus the experimental crystal structure. (A)** The original AF2 model had chain A (orange sticks) and chain B (green sticks) crossing over at residues N142-P144. Experimental $2F_0$-$F_c$ electron density maps are shown as a yellow mesh. **(B)** Colouring as for (A) demonstrates that rethreading of these chains matches the electron density in the experimental structure of B4GALNT1.

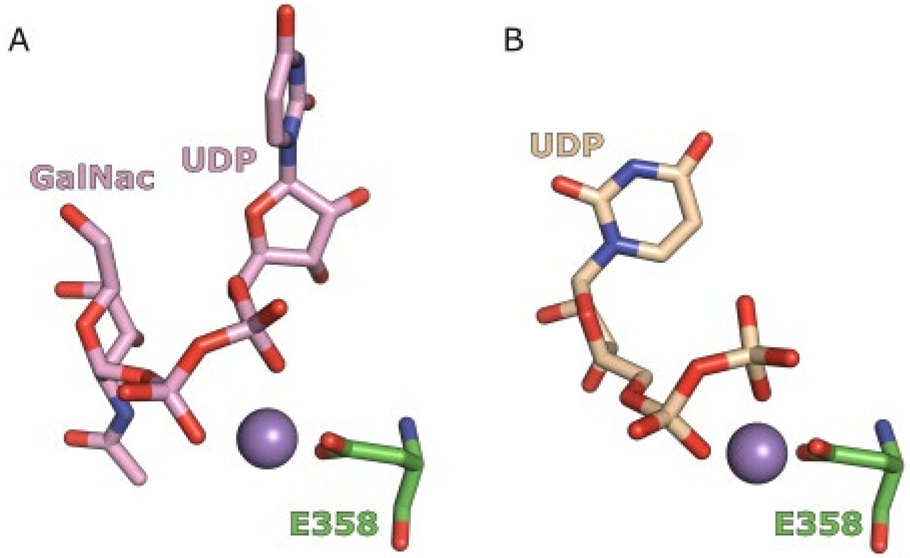


**Supplementary Figure S4.** Comparison of UDP substrate and product conformations in the B4GALNT1 active site. **(A)** The orientation of UDP-GalNac in the substrate complex is shown (pink sticks) relative to the Mn ion (purple) and residue E358 (green sticks) in the active site. **(B)** Orientation as for (A) but showing the UDP product (wheat sticks) in the active site. The UDP moiety is bound in an alternative conformation compared to the UDP in the UDP-GalNac-bound structure.

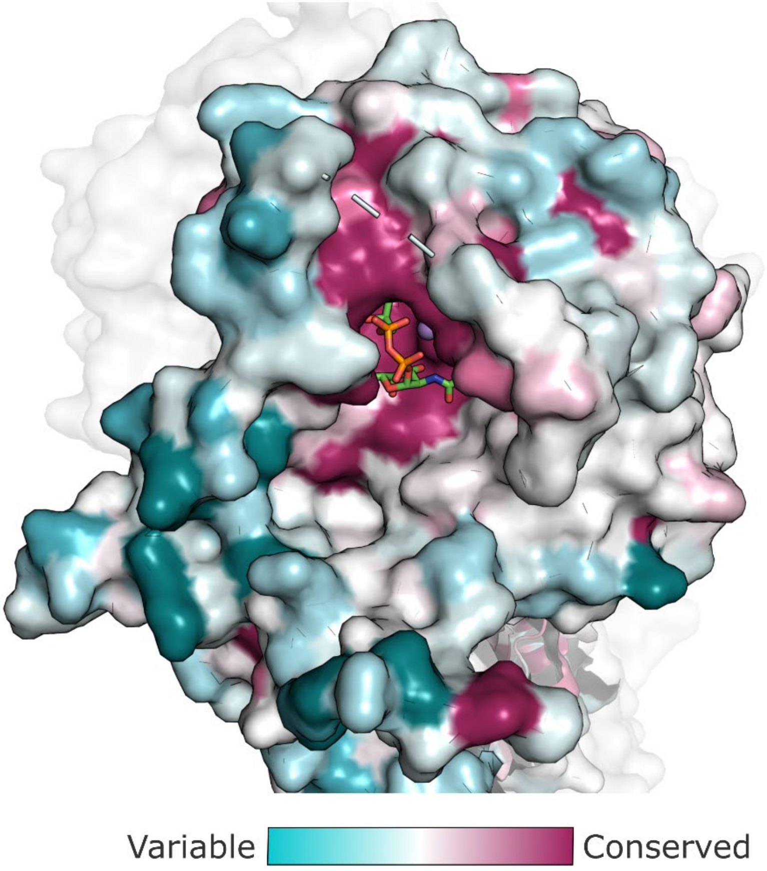


**Supplementary Figure S5.** Surface representation of the B4GALNT1 ligand binding site illustrating the UDP-GalNac (sticks) and showing conservation of residues across glycosyltransferase proteins. Sequence conservation was calculated and mapped onto the structure using Consurf [1] with colouring from highly variable (cyan) to highly conserved (magenta).

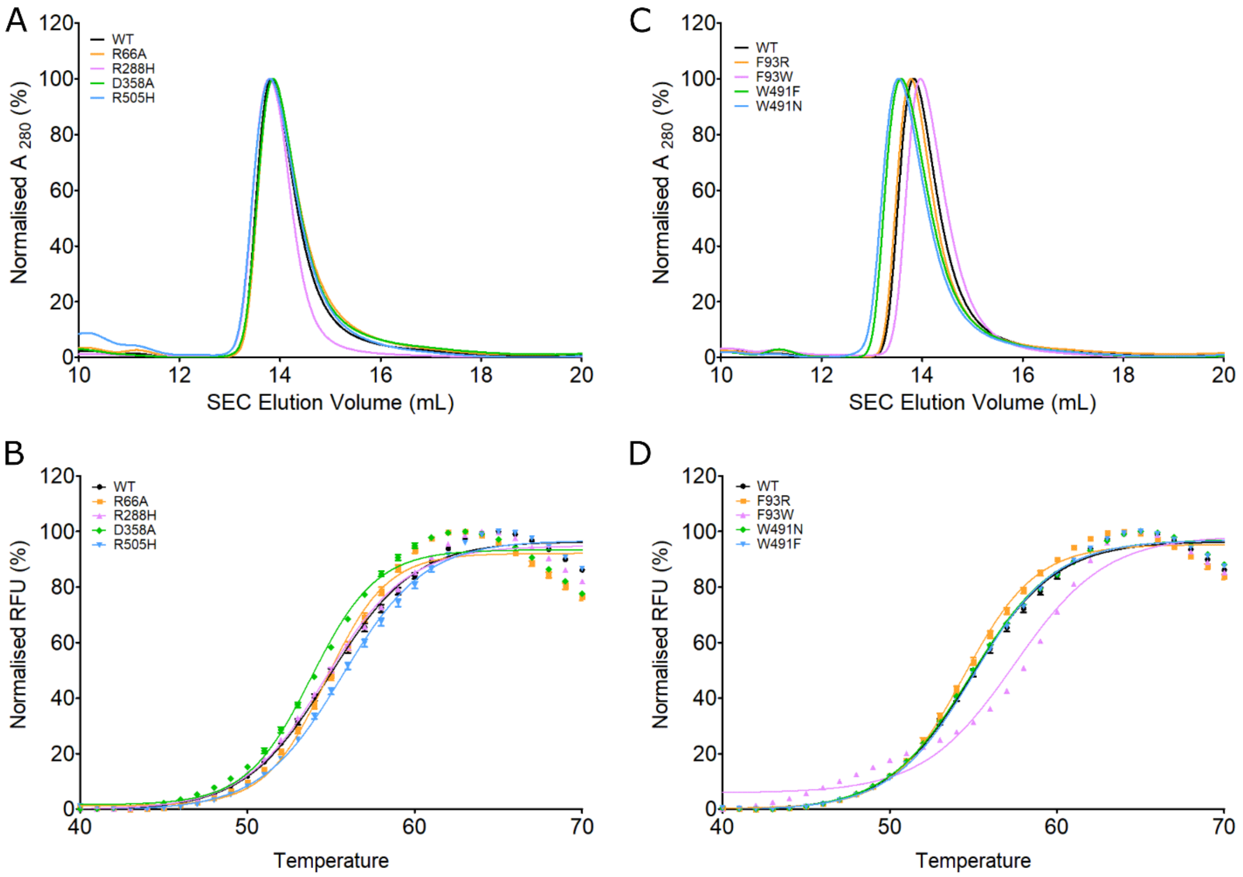


**Supplementary Figure S6.** Characterisation of B4GALNT1 mutants. **(A)** Elution profile following size-exclusion chromatography (SEC) for catalytic and disease mutants R66A, R288H, D358A and R505H. **(B)** Differential scanning fluorimetry (DSF) to monitor melting temperature as a measure of correct folding for the mutants as in (A). **(C)** SEC elution profiles for loop mutants F93R, F93W, W491F and W491N. **(D)** DSF melt curves for the loop mutants s in (C).

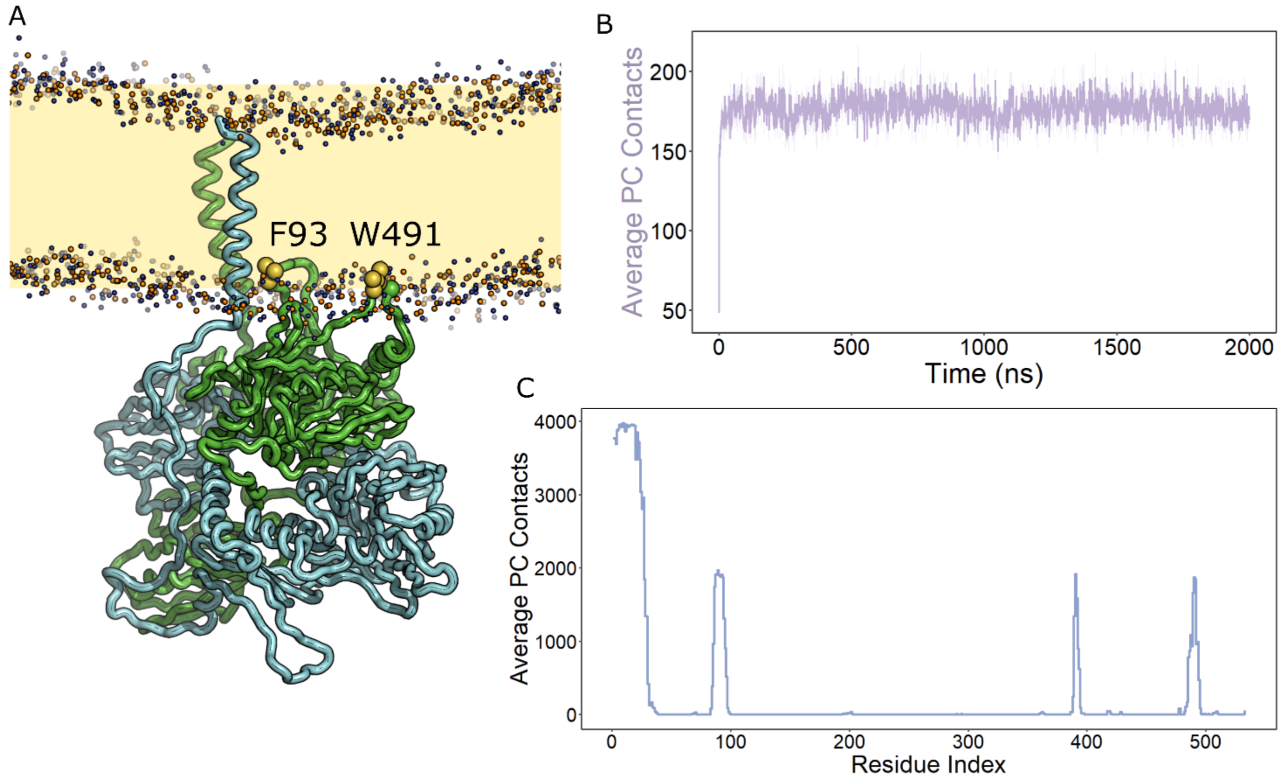


**Supplementary Figure S7.** Course-grain molecular dynamics simulations (MDS) for B4GALNT1 including the N-terminal transmembrane helix. **(A)** Illustration of a final pose of the full-length B4GALNT1 dimer in a PC membrane. As for the MDS with the luminal domain alone, the orientation of the lumenal domain consistently resulted in the insertion of the F93 and W491 loops of one chain of the dimer into the PC membrane. **(B)** Per residue analysis of the lipid contacts for the membrane-tethered B4GALNT1 reveal a similar pattern of luminal domain interactions with the membrane in addition to the transmembrane helix.

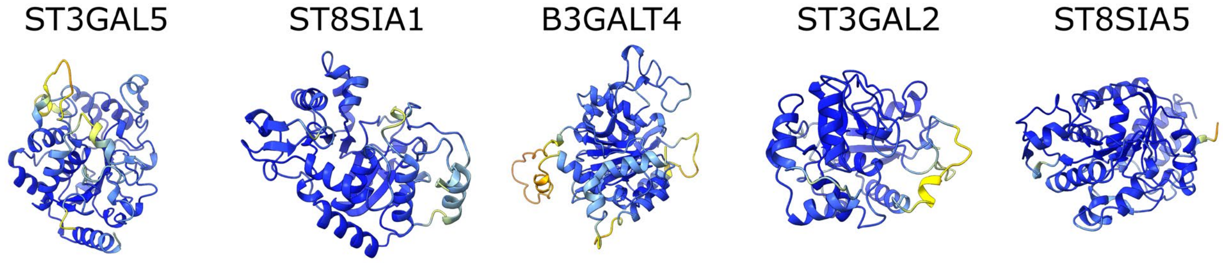


**Supplementary Figure S8.** Ribbon diagrams of AF2 predicted structures of the luminal domains of ganglioside synthetic enzymes coloured by pLDDT score. Colouring is from high confidence to low confidence prediction (blue through yellow to red).

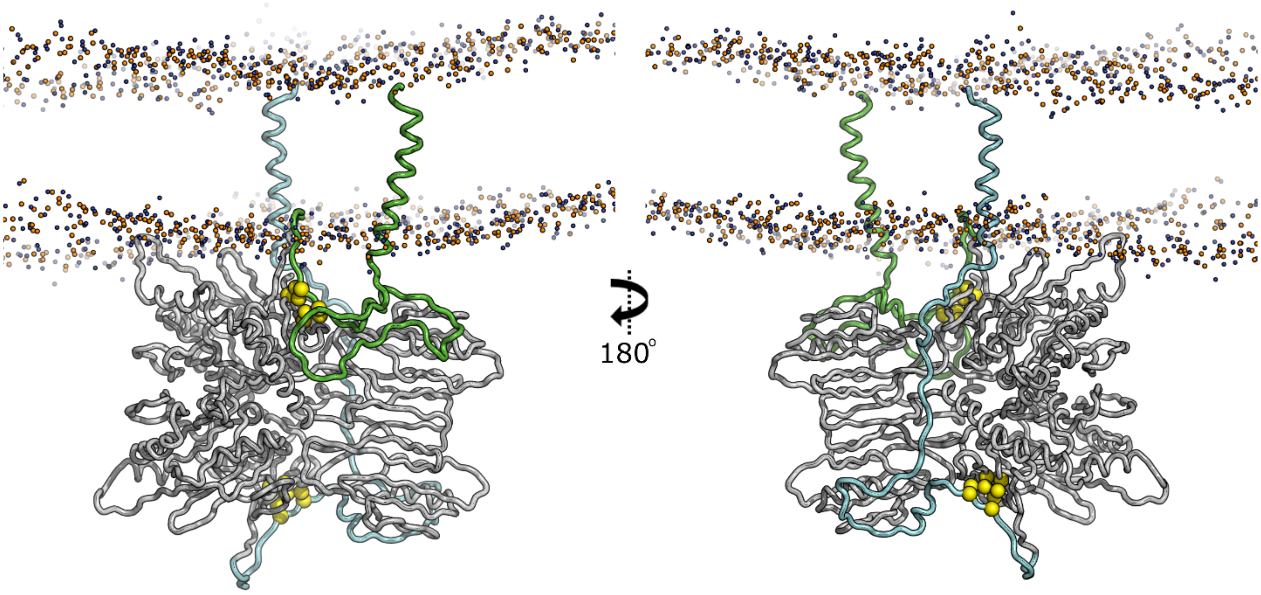


**Supplementary Figure S9.** Example poses of the full-length B4GALNT1 following MDS. For one molecule of the dimer the residues from the transmembrane helix to the N-terminal region of the active site (green) are not extended (left panel). This arrangement means that this region of the other molecule in the dimer (blue) is extended (right panel). This part of the molecule now spans from the membrane across the dimer and to the N-terminal region of the active site. The disulfide bonds that hold the N-terminal loop containing F93 are shown (yellow spheres).

**Supplementary Table 1.** B4GALNT1 missense mutations linked to HSP26

| Mutation | Predicted Impact Based on Structure | References |
|---|---|---|
| K284N | Catalytic: directly binds UDP-GalNac substrate | [2–5] |
| R300C | Misfolding: stabilises fold in non-catalytic domain | [2,3,5,6] |
| D433A | Destabilises dimer: bonds to H420 that pi-stacks with Y230 at dimer interface | [2–7] |
| P453R | Destabilises dimer: mediates a backbone turn at dimer interface | [2] |
| R472P | Destabilises dimer: forms H-bonds at dimer interface | [7] |
| S475F | Misfolding: buried, pocket can't fit bigger, hydrophobic F sidechain | [8] |
| R505H | Catalytic: directly binds UDP-GalNac substrate | [2–5] |
| R505C | Catalytic: directly binds UDP-GalNac substrate | [7] |
| R519P | Misfolding: Pro breaks helix and breaks H-bond | [2] |
